# GeneSurv: Multi-Gene Cancer Survival Analysis Tool

**DOI:** 10.1101/2024.03.17.583882

**Authors:** Zikang Xu, Jamin Wu, Sung-young Shin, Lan K. Nguyen

## Abstract

**Summary:** Traditional cancer survival analysis tools often restrict their analysis to single genes, thereby limiting their analytical scope. GeneSurv, a novel Web-Based Multi-Gene Cancer Survival Analysis Tool, addresses this limitation by enabling multi-gene analyses that correlates the expression of multiple genes, without restriction on their number, in various combinations, with patient survival outcomes. GeneSurv offers dynamic functionalities and an intuitive interface for flexible and precise adjustments of gene expression thresholds. Among its demonstrated applications, GeneSurv identified a significant association between high expression of CCND1 and CDK4 with poor breast cancer prognosis, and related the combined expression of five genes to patient survival, showcasing its capacity to unravel the complexities of genetic interactions and their impact on patient outcomes.

**Conclusion:** Available with user-friendly tutorials at https://can-sat.erc.monash.edu/, GeneSurv is an invaluable addition to the existing suite of bioinformatic tools, significantly enhancing the investigation of cancer prognosis.

## Introduction

The accurate prediction of cancer prognosis is a critical objective in oncological research, as it directly informs treatment decisions and patient management strategies. By integrating gene expression data with clinical outcomes, researchers can uncover the prognostic significance of specific genes, thereby identifying potential biomarkers for cancer and identifying factors contributing to treatment resistance ^1^. However, despite the abundance of open cancer datasets, the processes of data collection, cleaning, and analysis remain significant challenges for both biologists and clinicians, hindering the efficient utilization of this wealth of information in advancing personalized medicine.

Survival analysis is a crucial statistical approach that connects patient characteristics and survival outcomes, giving rise to various bioinformatic tools, such as KMplot ^2^, SurvExpress ^1^, and OSpaad ^3^. Despite their contributions, these tools face critical limitations. Firstly, they predominantly analyze survival data based on single-gene expressions, which narrows the scope of understanding the complex interplay between multiple genes and its impact on patient survival. For instance, synergistic effects of multiple gene expressions could provide deeper insights into tumour progression or identify novel therapeutic targets. Secondly, the classification of patients into control and test groups typically relies on the median or quartiles of gene expression levels, which overlooks the nuanced roles that genes may play in cancer, thereby limiting the potential for more refined patient stratification. Thirdly, certain tools, such as OSpaad, are restricted to specific cancer types, mainly pancreatic cancer, and others like SurvExpress are not regularly updated, making them unable to analyze newer datasets. These issues underscore the need for a more sophisticated and regularly updated survival analysis tool that can accommodate the complexities of cancer research.

To address these challenges, we present the Web-Based Multi-Gene Cancer Survival Analysis Tool (GeneSurv), a platform that enables survival analysis across 94 publicly available cancer datasets. GeneSurv features a user-friendly interface suitable for users of varying expertise and integrates data from the cBioPortal API ^4^ to provide up-to-date information. A key feature of GeneSurv is its capacity for multi-gene analysis, recognizing the intricate interactions among multiple genes in cancer progression and offering a more detailed understanding of its genetic underpinnings. By enabling the analysis of gene combinations, GeneSurv not only aims to identify patterns in cancer progression that singlegene studies might miss, thereby expanding the scope of survival analysis in oncology, but its robust multi-gene analysis framework is also set to enhance our grasp of complex genetic interactions in cancer, potentially uncovering new biomarkers and therapeutic targets. We envision GeneSurv becoming an essential tool for the cancer research community, contributing to more comprehensive survival studies and fostering collaborative efforts.

## Methods

### Data acquisition and web interface implementation

GeneSurv integrates a comprehensive collection of 94 cancer datasets from cBioPortal ^4^, each including a minimum of 40 cases with recorded overall or disease-free survival data and corresponding mRNA expression profiles. Upon receiving an analysis request, GeneSurv’s backend dynamically fetches the gene expression data for selected genes from cBioPortal and integrates it with pre-cached clinical data. This process involves converting gene expression levels into percentiles relative to all cases in the selected study, thereby enabling a detailed and accurate categorization of cases into Test or Control groups based on user-specified gene expression thresholds.

The web interface of GeneSurv, developed using React for the frontend and Flask for the backend, prioritizes user accessibility and ease of use. It features an intuitive and responsive layout that simplifies navigation, gene selection, threshold setting, and analysis submission for users. The results, including survival curves and hazard ratios, are displayed directly in the user’s browser with interactive visualizations and options for in-depth data examination. This strategic fusion of an extensive data repository, real-time data retrieval, and a user-centric interface positions GeneSurv as a versatile and powerful tool for detailed analysis of survival outcomes across a various cancer types.

### Survival analysis

GeneSurv splits cases into Test or Control groups based on user selected genes and thresholds. For the survival analysis, GeneSurv employs the Kaplan-Meier model ^5^ and the Cox Proportional Hazards model ^6^, both of which are implemented in Python’s lifelines library ^7^, ensuring robust and reliable statistical analysis. The analysis provides a clear visualization of the correlation between gene expression and patient survival. A significant difference in survival is defined by a p-value less than 0.05. Additionally, a hazard ratio (HR) greater than 1 indicates increased risk, while an HR less than 1 suggests reduced risk.

## Results

### Key features

- **Comprehensive Cancer Dataset Access:** GeneSurv incorporates 94 diverse cancer datasets from cBioPortal, each comprising survival data and mRNA expression profiles. This aggregation enables broad analyses across different cancer types.
- **Multi-Gene Analysis Capability:** GeneSurv surpasses traditional single-gene analysis by employing a multi-gene analysis framework, without the limit on the number of genes. This is crucial for uncovering potential synergistic effects among genes and elucidating their collective impact on patient prognosis.
- **Multi-Gene Contrast Selection:** Besides conventional analyses that simply differentiate between high and low gene expression groups, GeneSurv enables comparisons between patients exhibiting high expression levels of certain genes (e.g. oncogenes) in tandem with low expression of others (e.g. tumour suppressors), and vice versa, offering a more sophisticated analysis of their combined influence on patient prognosis.
- **User-Friendly Web Interface:**
  - ***Interactive Analysis Tools:*** The platform offers dynamic features for gene selection, threshold adjustments, and case categorization, coupled with clear visualizations that display multiple analyses simultaneously, allowing straightforward cross-comparison and enhancing user interpretability.
  - ***‘Mirror Flip’ Feature:*** This innovative and convenient feature enhances group categorization in gene expression studies by enabling inverted threshold settings. This means when activated, a standard lower 30th percentile threshold reassigns the Control group to above the 70th percentile.
  - ***Reproducibility and Data Export:*** The tool promotes reproducibility by enabling the download of analysis configurations in JSON format, as well as clinical and gene expression data in CSV, and Kaplan Meier plots in PNG format.

### Interface and workflow

**Figure 1A** outlines GeneSurv’s workflow and user interface. Initially, users select a dataset from a dropdown menu, and upon selection, a dataset description and options for survival outcome and gene expression profiles appear. While default settings are provided, certain studies offer additional options, enabling users to select the most relevant mRNA expression profile for their research. Users then input at least one gene of interest, triggering the appearance of an adjustable slider for setting gene-specific criteria. The blue and grey regions indicate the Test and Control criteria for each gene, respectively, with both groups needing to meet the set criteria for all genes. For instance, **Figure 1A** shows control group patients with ERBB2, CCND1 and CDK6 gene expression in the bottom 30%, 70%, and 50%, respectively. The mirror feature, indicated by a greyed-out button when activated, alters the ERBB2 expression analysis in the test group to the top 30%. In contrast, CCND1 and CDK6, without the mirror feature activated, are displayed in the top 30% and 50%, highlighting the feature’s role in data representation. Users should exercise caution when selecting three or more genes or using overly strict criteria to avoid a low case count that could hinder meaningful analysis.

**Fig. 1.**
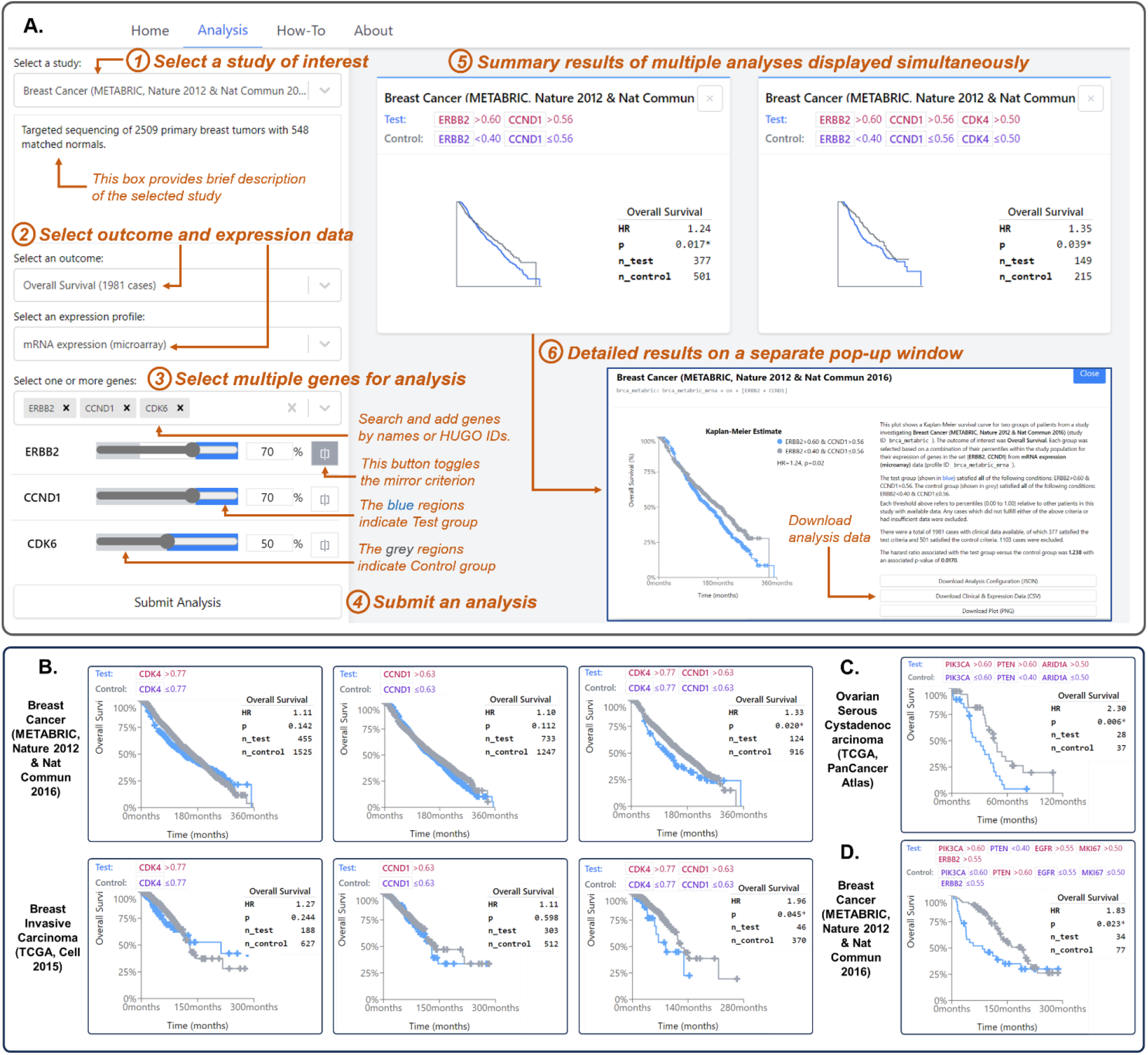
(A) Step-by-step usage of MGSAT. 1)Select a study from the dropdown list. 2) Select the survival outcome and gene expression profile. 3) Select the gene(s) you wish to combine and their levels. 4) Click button to submit an analysis and it will be added to the right. 5) Click on the summary to open the analysis in more detail. 6) Click these buttons to download data related to the analysis. **(B) Survival association with CDK4 and CCND1 gene expression across two datasets**. Combined high expression of CCND1 and CDK4 in both datasets is significantly associated with poorer breast cancer prognosis (P < 0.05, HR > 1), contrasting with individual gene overexpression which showed no significant survival impact. **(C) Association of PIK3CA, PTEN, and ARID1A Expression with Ovarian Cancer Prognosis. (D) Prognostic Implications of Combined Gene Expression in Breast Cancer**.

After finalizing gene selections and thresholds, the analysis is initiated with the ‘Submit Analysis’ button, and the results start to load on the right side. Once completed, a summary card appears, clickable for a detailed view including a textual description and a Kaplan Meier plot. For reproducibility, users can download the analysis configuration in JSON format, encompassing all analysis details such as selected study, outcomes, expression profiles, and gene thresholds. Additionally, a CSV file containing clinical and gene expression data (with genes identified by ENTREZ ID) and a PNG file of the Kaplan Meier plot are available for download. Storing these files is recommended for future analysis replication.

To help users navigate and effectively utilize the platform, a comprehensive ‘How-To’ guide is available on the website (https://can-sat.erc.monash.edu/). This guide provides step-by-step instructions and tips for maximizing the tool’s potential, ensuring users of varying expertise levels can conduct their analyses with ease and confidence.

### Case Studies

To showcase GeneSurv’s utility, here we demonstrated its application through case studies on breast cancer and ovarian serous cystadenocarcinoma, revealing new insights into the complex interplay of gene expressions and their impact on cancer prognosis.

### Combined high expression of CCND1 and CDK4 is linked to poor prognosis in breast cancer

Cyclin D1 (CCND1) and its binding partner CDK4 are transcriptional regulators that control cell proliferation and migration^8^. Mutations in CCND1 and CDK4 have been linked to an increased risk of breast cancer^9^; however the relationship between their gene expression and breast cancer prognosis remains unclear. Filho et al.^10^ observed no significant association between CCND1 expression and overall survival in basal-like breast cancer cells. Conversely, Valla et al.^11^ associated higher CCND1 copy numbers with more aggressive tumor characteristics, but not directly with prognosis. Peurala et al.^12^ reported no significant association between CDK4 gene expression with prognosis. Overall, the combined effect of CCND1 and CDK4 expression on patient prognosis has yet to be explored.

We utilized GeneSurv to analyze the relationship between CCND1 and CDK4 expression and patient survival using the METABRIC breast cancer^13–15^ and TCGA Breast Invasive Carcinoma^16^ datasets. Figure 1B shows our initial assessment of their individual expression levels, which found no significant association with overall patient survival, consistent with previous studies. However, GeneSurv’s multi-gene analysis revealed a significant link between their combined high expression and poor prognosis in both datasets. This suggests a synergistic effect when both genes are concurrently upregulated, offering directions for future research.

### Combined high expression of PIK3CA, PTEN, and ARID1A linked to poor prognosis in ovarian cancer

Here, we used GeneSurv to analyse the Ovarian Serous Cystadenocarci-noma dataset from the TCGA PanCancer Atlas, focusing on the impact of PIK3CA, PTEN, and ARID1A gene expression and patient prognosis. PIK3CA is known for its role in the PI3K/AKT/mTOR signaling pathway and tumorigenesis in ovarian cancer^17^ while PTEN is a tumor suppressor in the same pathway^18^. Studies suggest that high PTEN expression can sensitize ovarian cancer cells to chemotherapy, potentially through a p53-mediated apoptotic cascade, independent of the PI3K/AKT pathway^19^. On the other hand, low PTEN expression correlates with enhanced tumorigenic activity due to reduced PI3K/AKT pathway suppression^20^. To reflect its complex role in cancer, PTEN gene expression was examined at both high (>0.6) and low (<0.4) expression levels using GeneSurv’s mirrored criteria feature. ARID1A contributes to chromatin remodeling and, when altered, can affect the expression of multiple genes, thereby influencing carcinogenesis, often in conjunction with the PI3K/AKT pathway^21^.

Our analysis revealed a significant association between high expressions of PIK3CA, PTEN, and ARID1A (>0.6, >0.6, and >0.5, respectively) and poorer overall survival, with a notable increase in hazard ratio (HR = 2.3, P-value = 0.006) in the Test group compared to the Control group with lower expressions of these genes (Fig. 1C). This finding offers deeper insights into how these multiple genes collectively influence ovarian cancer progression and patient outcomes, thereby enhancing our understanding of gene interaction dynamics in cancer.

### High PIK3CA, EGFR, MKI67, ERBB2 coupled with low PTEN expression correlate with poor prognosis in breast cancer

In this case study, we leveraged the unique multi-gene feature of GeneSurv tool to explore the link between the expression of five key genes - PIK3CA, PTEN, EGFR, MKI67, and ERBB2 – and prognosis in breast cancer. The rationale for selecting these specific genes is based on their established individual roles in breast cancer progression, yet the impact of their interaction in prognosis is unclear. PIK3CA alterations, which activates the PI3K/AKT/mTOR pathway, are common in breast cancer^22^. PTEN acts as a key inhibitor of this pathway, with its expression levels influencing tumour behavior^23^. EGFR overexpression is linked to increased tumour aggressiveness and poorer prognosis^24^. MKI67 serves as a marker of proliferation marker, with high levels indicating rapid tumor growth and negative prognosis^25^. ERBB2 (HER2) overexpression is a well-known prognostic factor in certain breast cancer subtypes, significantly affecting patient survival^26^.

Using the METABRIC dataset^13–15^, our analysis revealed a significant increase in hazard ratio (HR = 1.83, P-value = 0.023) in the Test group (PIK3CA >0.60, PTEN <0.40, EGFR >0.55, MK167 >0.50, ERBB2 >0.55) compared to the Control group (Fig. 1D). Notably, omitting any of the genes PIK3CA, PTEN, MKI67, or ERBB2 nullified the statistical significance of the results, while excluding EGFR reduced the significance, suggesting they act synergistically to influence patient prognosis. These findings highlight GeneSurv’s ability for complex multi-gene analysis, providing new insights into the impact of genetic interactions in cancer.

## Conclusion

GeneSurv surpasses the constraints of current survival analysis tools by facilitating detailed multi-gene analyses. Its dynamic capabilities, intuitive interface, and the novel ‘Mirror Flip’ feature for precise gene expression threshold adjustments allow researchers to unravel the complexities of genetic interplay and their effects on patient outcomes. With these advancements, GeneSurv is poised to become an invaluable addition to the existing suite of bioinformatic tools, significantly enhancing the investigation of cancer prognosis.

## Acknowledgments

We are grateful to Monash University for providing the computing resources necessary to host GeneSurv’s webpage and execute the analyses.

